# Peripapillary Perfused Capillary Density in True Versus Pseudoexfoliation Syndrome: An OCTA Study

**DOI:** 10.1101/2020.09.01.277293

**Authors:** Phantaraporn Tangtammaruk, Purit Petpiroon, Wasu Supakonatanasan, Chaiwat Teekhasaenee, Yanin Suwan

**Author notes:** Correspondence author, Department of Ophthalmology, Ramathibodi Hospital, Mahidol University, 270 Rama VI, Thung Phaya Thai, Ratchathewi, Bangkok, Thailand, 10400.

## Abstract

**Purpose:** To compare peripapillary perfused capillary density (PCD) among eyes with true exfoliation syndrome (TEX), eyes with pseudoexfoliation syndrome (PEX), and healthy control eyes.

**Materials and Methods:** In this observational cross-sectional study, eyes with and without TEX or PEX were assessed by optical coherence tomography angiography (OCTA) imaging. Bilateral OCTA images (4.5 · 4.5 mm^2^) centered at the optic nerve head were obtained using a commercial spectral domain OCTA system. Optic nerve head perfusion was quantified using the split-spectrum amplitude decorrelation angiography algorithm. Categorical and continuous variables were compared using the chi-squared test and one-way analysis of variance, respectively. The generalized estimating equation was used to adjust for confounding factors and determine inter-ocular associations.

**Results:** We enrolled 39 eyes with TEX, 31 eyes with PEX, and 32 control eyes. There were no significant differences among the three groups regarding age, intraocular pressure, cup-to-disc ratio, blood pressure, or axial length (all p>0.05). There were significant differences in global PCD among the three groups (p=0.01). There were significant differences in annular PCD between the TEX and PEX groups (p=0.027).

**Conclusions:** While both global and annular PCDs did not differ between the TEX and control groups, greater loss of annular PCD in the PEX group than in the TEX and control groups suggests more pronounced microvascular disturbance in PEX.

**Synopsis/Precis:** Greater microvascular attenuation in PEX compared with TEX and normal control measured by OCTA.

## Introduction

True exfoliation syndrome (TEX) is a disease characterized by delamination of anterior lens capsule, which was first reported in 1922 by Elsching.^1,2^ Occupational exposures by steelworkers, blacksmiths, glassblowers, and bakers may increase the risk of TEX.^3^ Several studies have shown that many patients exhibit idiopathic forms of TEX.^2,3^ Nonetheless, vascular insufficiency has not been studied with respect to TEX pathogenesis.

Pseudoexfoliation syndrome (PEX) is an age-related systemic disorder characterized by the production and accumulation of pseudoexfoliation material. PEX is one of the most common causes of secondary open-angle glaucoma.^4^ Several studies have shown an association between PEX and increased systemic vasculopathy, as well as between PEX and ocular vasculopathy.^5-15^ It has been proposed that pericellular accumulation of pseudoexfoliation material may disrupt the normal cellular basement membrane, thereby leading to endothelial dysfunction.^16^ Iris vasculopathy, endothelial basement membrane abnormalities, and lumen obliteration have been described in patients with PEX.^5–7^ In addition to altered intraocular pressure, ocular blood flow disturbance constitutes evidence of the pathogenesis of pseudoexfoliation glaucoma progression.^5^

Vascular insufficiency is a notable risk factor for glaucoma pathogenesis. Various methods have been used to measure optic nerve head perfusion in both glaucoma and PEX.^5^ Fluorescein angiography has been used to measure vascular blood flow; however, it is both invasive and inaccurate for measurement of optic nerve head perfusion. Laser speckle flowgraphy and Doppler imaging are operator-dependent and do not provide adequate visualization. Furthermore, these methods do not distinguish some small complex vessels in the optic disc.^17^

Optical coherence tomography angiography (OCTA) is a noninvasive imaging modality that combines optical coherence tomography (OCT) reconstruction with motion contrast processing to identify retinal microvasculature.^18^ Creation of a retinal blood flow map is performed using the split-spectrum amplitude decorrelation angiography algorithm, which compares the decorrelation signals among sequential OCT B-scans taken at the same cross-sectional level.^8^

TEX is presumed to be a subtype of PEX. Previous OCTA studies of peripapillary perfused capillary density (PCD) in patients with PEX, pseudoexfoliation glaucoma, or primary open-angle glaucoma provided evidence of microvascular disturbances.^8,19^ However, to the best of our knowledge, there has been no evidence of vasculopathy in patients with TEX. The purpose of this study was to compare PCD among eyes with TEX, eyes with PEX, and healthy control eyes.

## Materials and methods

This cross-sectional study was approved by the Center of Ethical Reinforcement for Human Research at Mahidol University. Written informed consent was obtained from all participants before enrollment, and the study protocol adhered to the tenets of the Health Insurance Portability and Accountability Act and the Declaration of Helsinki.

### Participants

Patients who were diagnosed with TEX or PEX at the Department of Ophthalmology in Ramathibodi Hospital were recruited between March 2018 and March 2019. The inclusion criteria for all patients were Snellen best-corrected visual acuity ≥ 20/40 and age > 20 years. The diagnosis of TEX was based on characteristic anterior lens capsule delamination, intraoperative double-ring sign, and pigment deposition on delaminated membrane. PEX was diagnosed by the presence of pseudoexfoliation material on slit-lamp biomicroscopy with normal optic disc and no glaucomatous damage detected by OCT or perimetry.

Inclusion criteria for healthy control eyes were defined as follows: absence of retinal pathology or glaucoma, intraocular pressure ≤ 21 mmHg, no chronic ocular or systemic corticosteroid use, open anterior chamber angle on gonioscopy, absence of pseudoexfoliation material or delaminated lens capsule, normal-appearing optic nerve head, and no glaucomatous damage detected by OCT or perimetry. Exclusion criteria for all participants were eyes with a history of ocular surgery except uncomplicated cataract surgery, any ocular diseases other than cataract, history on systemic steroid usage, diabetes mellitus, and cardiovascular disease other than treated systemic hypertension. All participants underwent a standard ophthalmic examination, fully dilated ophthalmoscopy, medical history review, systolic and diastolic blood pressure measurements, best-corrected visual acuity assessment, Goldmann applanation tonometry, and A-scan biometry. Automated perimetry was performed on the Humphrey Visual Field Analyzer (Humphrey Instruments Model Model 740; Carl Zeiss Meditec, Dublin, CA, USA) using the 24-2 Swedish Interactive Testing Algorithm standard.

### Image acquisition and scanning protocol

All eyes were imaged using a 3.45-mm circle OCT scan centered at the optic nerve head for circumpapillary retinal nerve fiber layer (RNFL) thickness analysis (Spectralis OCT Version 6.0.11.0 + Glaucoma Module Premium Edition Software; Heidelberg Engineering, GmbH, Dossenheim, Germany). Scans with signal strength <7/10 were excluded.

A spectral domain OCT system (AngioVue Software Version 2018.1.0.33, Optovue, Fremont, CA, USA) was used with a single imaging session consisting of two volumetric raster scans (one vertical and one horizontal) centered on the optic nerve head and covering an area 4.5. 4.5 mm. Perfused vessel imaging was performed using the split-spectrum amplitude decorrelation angiography algorithm. The vessel located between the internal limiting membrane and the posterior boundary of the RNFL was identified as the radial peripapillary capillary. Images with signal strength index ≤ 40 and those with eye motion artifacts were excluded (Figure 1).

**Figure 1.**
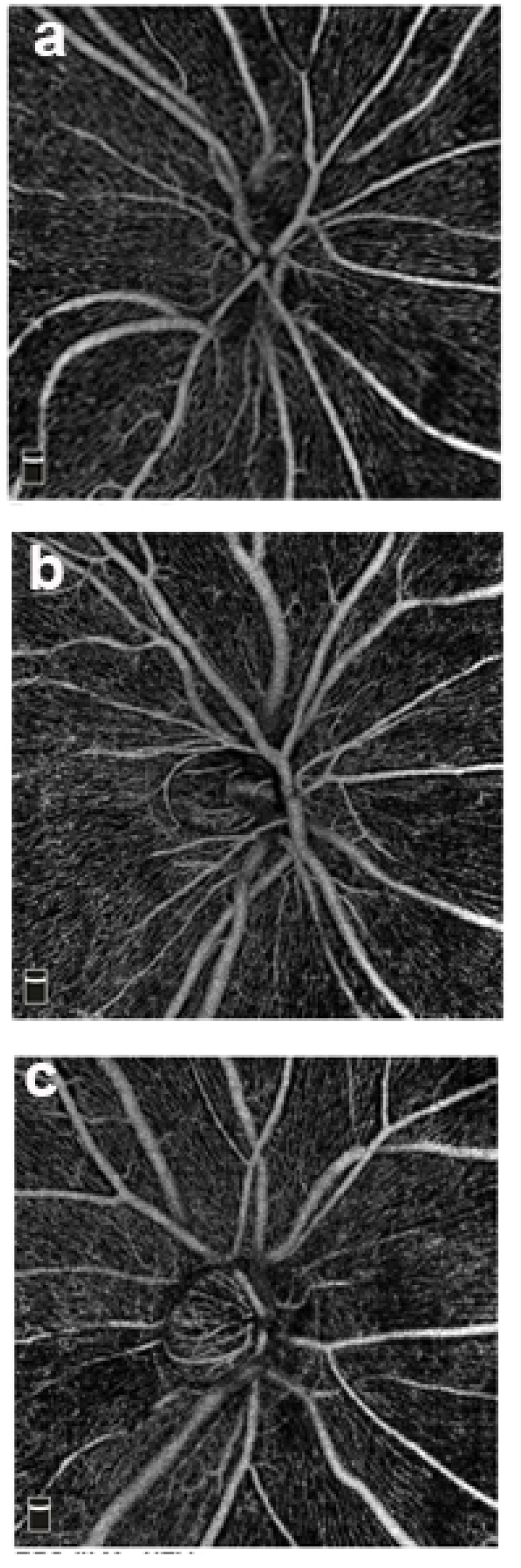
Optical Coherence Tomography Angiography (OCTA) Images of the Three Groups. OCTA images in gray scale. a: control, b: true exfoliation syndrome, c: pseudoexfoliation syndrome

### OCTA image analysis

OCTA images were analyzed using built-in Optovue software^18, 20^ with the large blood vessel removal algorithm.

### Statistical analysis

In patients with bilateral pathology or healthy eyes (i.e., both eyes with TEX or PEX, or both healthy controls), both eyes were included for analysis. In patients with unilateral pathology, the eye with disease was selected for analysis. The normality of the numerical data distribution was assessed using the Shapiro–Wilk test. Descriptive statistics were calculated as the mean and standard deviation for normally distributed variables. Categorical and continuous variables were compared using the chi-squared test and one-way analysis of variance, respectively; a Bonferroni post hoc test was used to assess the differences between pairs of groups following analysis of variance. The generalized estimating equation was used to determine interocular associations, adjusted for age, sex, axial length, and blood pressure. Differences with p<0.05 were considered statistically significant. All statistical analyses were performed using Stata, version 14 (STATA Corp., College Station, TX, USA).

## Results

Initially, there were 48 eyes with TEX, 34 eyes with PEX, and 32 control eyes in this study. After excluding 12 eyes with low signal strength scans, the eyes included in this study were 39 with TEX (mean patient age [standard deviation], 74.13 [7.34] years), 31 with PEX (mean patient age [standard deviation], 73.45 [9.64] years) and 32 controls (mean patient age [standard deviation], 71.38 [5.12] years). Demographic characteristics among the three groups are demonstrated in Table 1. There were no significant differences in axial length, cup-to-disc ratio, or blood pressure among the three groups (all p>0.05). There was a significant reduction in global circumpapillary RNFL in the PEX group (p=0.018). There were significant differences in sectoral circumpapillary RNFL thicknesses among the three groups, including marked differences in the inferior and temporal sectors (p<0.001).

**Table 1.** Clinical and demographic data of TEX, PEX and control groups.

|  | TEX<br>(N=39) | PEX<br>(N=31) | Control<br>(N=32) | P-value |
| --- | --- | --- | --- | --- |
| Age(y) | 74.13 ± 7.34 | 73.45 ± 9.06 | 71.38 ± 5.12 | 0.276 |
| Gender (Female %) | 27 (69.23) | 12 (38.71) | 20 (58.72) | 0.152‡ |
| IOP | 12.56 ± 2.77 | 12.65 ± 2.18 | 12.97 ± 2.74 | 0.759 |
| AXL(mm.) | 23.16 ± 1.11 | 23.33 ± 0.81 | 23.33 ± 1.08 | 0.738 |
| CD(ratio) | 0.37 ± 0.12 | 0.37 ± 0.08 | 0.34 ± 0.61 | 0.232 |
| SBP(mmHg) | 126.92 ± 15.29 | 129.97 ± 12.14 | 124.19 ± 14.15 | 0.268 |
| DBP(mmHg) | 72.33 ± 6.79 | 71.84 ± 7.46 | 73.69 ± 5.47 | 0.515 |
| **OCT** |  |  |  |  |
| Global(um) | 100.08 ± 9.61 | 93.55 ± 13.84 | 100.97 ± 10.10 | 0.018* |
| Superior(um) | 117.59 ± 14.10 | 112.71 ± 19.38 | 132.16 ± 22.99 | <0.001** |
| Inferior(um) | 126.13 ± 19.88 | 116.58 ± 26.03 | 128.28 ± 18.60 | 0.076 |
| Nasal(um) | 83.31 ± 11.55 | 81.61 ± 26.97 | 93.25 ± 15.53 | 0.029* |
| Temporal(um) | 76.46 ± 8.77 | 66.19 ± 8.84 | 75.75 ± 11.52 | <0.001** |
| **PCD** |  |  |  |  |
| Global PCD (%) | 47.95 ± 4.00 | 45.64 ± 5.12 | 48.74 ± 3.11 | 0.010* |
| Annular PCD (%) | 52.12 ± 6.18 | 49.28 ± 6.09 | 51.12 ± 3.80 | 0.104 |
| Superior PCD (%) | 53.28 ± 8.58 | 49.45 ± 7.27 | 51.59 ± 3.97 | 0.081 |
| Inferior PCD (%) | 54.75 ± 7.91 | 48.42 ± 9.58 | 51.34 ± 4.62 | 0.003** |
| Nasal PCD (%) | 50.26 ± 7.63 | 48.32 ± 9.42 | 49.81 ± 6.52 | 0.581 |
| Temporal PCD (%) | 53.00 ± 5.73 | 46.87 ± 8.87 | 51.00 ± 5.14 | 0.001** |
| Baseline clinical data reported as mean ± standard deviation |  |  |  |  |
| P-value among 3 groups (1-way ANOVA) |  |  |  |  |
| ‡P-value among 3 groups (Chi-square test) |  |  |  |  |
| Level of significant *95%, **99% |  |  |  |  |
| IOP= intraocular pressure, AXL= axial length, CD=cup-disc ratio, SBP= systolic blood pressure, DBP= diastolic blood pressure, OCT= Optical Coherence tomography, PCD= Perfused capillary density |  |  |  |  |

With respect to OCTA analysis, global PCD demonstrated significant differences among the three groups (p=0.01), while annular PCD did not (p=0.104). Differences in global and annular PCDs are shown in Tables 2 and 3. Greater losses of temporal and inferior PCDs were observed among the three groups (p=0.001 and p=0.003, respectively; Table 1). Pairwise comparison showed that both global and annular PCDs did not significantly differ between the TEX and control groups (p=0.564 and p=0.555, respectively; Table 2). Conversely, both global and annular PCDs were significantly lower in the PEX group than in the control group (p=0.011 and p=0.017, respectively; Table 2). Finally, annular PCD significantly differed between PEX and TEX groups (p=0.027; Table 3).

**Table 2.** Differences in globular, annular PCD among TEX and PEX comparing with control group.

| Group | Mean PCD Difference (%) |  |  |  |  |  |
| --- | --- | --- | --- | --- | --- | --- |
|  | Global | P-value | 95% Confidence Interval | Annular | P-value | 95% Confidence Interval |
| Control |  |  | NA |  |  | NA |
| TEX | -0.62 | 0.564 | (-2.723, 1.485) | 0.85 | 0.535 | (-0.197, 3.675) |
| PEX | -2.75 | 0.011 | (-4.856, -0.643) | -2.21 | 0.017 | (-4.036, -0.391) |

Analyzed by generalized estimation equation
PCD= Perfused capillary density, TEX= True exfoliation syndrome, PEX= Pseudoexfoliation syndrome

**Table 3.** Differences in globular, annular PCD between TEX and PEX.

| Group | Mean PCD Difference (%) (TEX and control group N=70) |  |  |  |  |  |
| --- | --- | --- | --- | --- | --- | --- |
|  | Global | P-value | 95% Confidence Interval | Annular | P-value | 95% Confidence Interval |
| PEX |  |  | NA |  |  | NA |
| TEX | 1.59 | 0.086 | (-0.226, 3.416) | 2.81* | 0.027 | (0.321, 5.302) |

Analyzed by generalized estimation equation
Level of significant \*95%,
PCD= Perfused capillary density, TEX= True exfoliation syndrome, PEX= Pseudoexfoliation syndrome

## Discussion

Our study revealed significant differences in PCD among the three groups. There was a significant reduction of PCD in eyes with PEX, compared with control eyes. Eyes with PEX showed reduced capillary perfusion, compared with other groups, in terms of global and annular scans. However, PCD did not significantly differ between TEX and control groups.

Multivariable analysis demonstrated a significant reduction in PCD in the PEX group, compared with the TEX group. This finding is consistent with the results of a previous study by Suwan et al.,^8^ which showed greater reduction of PCD in eyes with PEX than in eyes with primary open-angle glaucoma or control eyes. There is considerable evidence to suggest that PEX is associated with vascular pathology.^21, 22^ The mechanism of glaucomatous optic neuropathy in eyes with PEX-related glaucoma is due to mechanical damage by elevated intraocular pressure, which results in death of retinal ganglion cells^23^ and impairment of the retrograde axon;^24^ the neuropathy is also due to the deposition of PEX material within vasculature, which increases vascular resistance and reduces blood flow. This vasculopathy may produce synergistic effect on the natural course of PEX-related glaucoma, which may result in greater intraocular pressure fluctuation, more severe clinical manifestations, and greater recalcitrance, compared with primary open-angle glaucoma. Shawn et al. found lower macula vessel density in eyes with pseudoexfoliation glaucoma than in eyes with primary open-angle glaucoma, which constitutes evidence for vasculopathy in eyes with PEX.^25^ Conversely, there is no evidence to support reduction of optic nerve head blood flow in eyes with TEX.

Our study showed that eyes with PEX had significant reduction in global circumpapillary RNFL, compared with healthy controls. This is consistent with the findings of a previous study by Yuksel et al., which found that PEX may be associated with a thinner RNFL, compared with normal eyes.^26^ Laminar elastosis pathogenesis might explain thinning RNFL in eyes with PEX.^27^ Gokhan et al. also reported that thinning of the RNFL was associated with the progression of PEX to pseudoexfoliation glaucoma.^28^

Our study suggests that vascular risk factors are not present in TEX. The presumed pathogenesis of TEX is idiopathic or heat-induced structural alteration of the lens capsule.^2, 3^ Physiologic iris movement and obstructed aqueous flow in the iris–lens channel have been proposed as contributing factors for the progression of detached membrane in eyes with TEX. These suggest a local, rather than systemic, insult to ocular tissue in eyes with TEX; in contrast, PEX is known as an ocular manifestation of a systemic condition that involves accumulation of abnormal fibrillary material.

There were some limitations in our study. In addition to the small sample size, the cross-sectional study design limited the statistical power of the findings and prevented follow-up regarding disease progression. Moreover, our study included eyes with any stage of TEX, which may have led to differences in PCDs that were recorded. Future studies should include more precise staging of TEX to avoid bias in the PCD results. In our study, assessment of control eyes included maximal mydriasis with slit-lamp examination to exclude eyes with early-stage TEX.

To the best of our knowledge, this is the first quantitative comparison of the degree of microvascular disturbance among eyes with PEX, eyes with TEX, and healthy control eyes. OCTA imaging might be beneficial for detection of vessel density loss. Our findings suggested greater microvascular attenuation in eyes with PEX than in eyes with TEX or control eyes, which implies that PEX and TEX are separate diseases, rather than two subtypes of a single disease. A longitudinal study with a larger sample size is warranted to confirm these findings.

